# Comparative roles of caudate and putamen in the serial order of behavior: Effects of striatal glutamate receptor blockade on variable versus fixed spatial self-ordered sequencing in marmosets

**DOI:** 10.1101/2023.12.20.572571

**Authors:** Stacey Anne Gould, Amy Hodgson, Hannah F. Clarke, Trevor W. Robbins, Angela C. Roberts

**Affiliations:** Department of Physiology, Development and Neuroscience, University of Cambridge, Downing Street, Cambridge, CB2 3DY, UK; Department of Psychology, University of Cambridge, Downing Street, Cambridge, CB2 3EB, UK

**Author notes:** Correspondence to: Stacey Anne Gould, Department of Physiology, Development and Neuroscience, University of Cambridge, Downing Street, Cambridge, CB2 3DY, UK. Joint senior authors (Trevor W. Robbins, Angela C. Roberts).

## Abstract

Self-ordered sequencing is an important executive function involving planning and executing a series of steps to achieve goal-directed outcomes. Lateral frontal cortex is implicated in this behavior, but downstream striatal outputs remain relatively unexplored. We trained marmosets on a three-stimulus self-ordered spatial sequencing task using a touch-sensitive screen to explore the role of caudate nucleus and putamen in random and fixed response arrays. By transiently blocking glutamatergic inputs to these regions, using intra-striatal CNQX microinfusions, we demonstrate that caudate and putamen are both required for, but contribute differently to, flexible and fixed sequencing. CNQX into either caudate or putamen impaired variable array accuracy, and infusions into both simultaneously elicited a greater impairment. We demonstrate that continuous perseverative errors in the variable array were caused by putamen infusions, likely due to interference with the putamen’s established role in monitoring motor feedback. Caudate infusions, on the other hand caused recurrent perseveration, with deficits possibly reflecting interference with the caudate’s established role in spatial working memory and goal-directed planning. In contrast to the variable array, whilst both caudate and putamen are needed for fixed array responding, combined effects were not additive, suggesting possible competing roles. Infusions in either region led to continuous perseveration when infused individually, but not when infused simultaneously. Caudate infusions did not cause recurrent perseveration in the fixed array; instead, this was caused by putamen infusions. The results overall are consistent with a role of caudate in planning and flexible responding, but putamen in more rigid habitual or automatic responding.

**Significance Statement:** This investigation employing local intra-striatal infusions into caudate nucleus and/or putamen of the New World marmoset reveals important roles for these regions in variable and fixed spatial self-ordered sequencing. Here, we directly implicate subcortical output regions of the lateral prefrontal cortex in self-ordered sequencing behavior. The ability to self-order sequences, as well as more broadly to plan, organize information, and respond flexibly, is impaired in many neurological diseases and psychiatric disorders. By understanding the basic neural circuitry underlying these cognitive abilities, we may better understand how to rectify them in people with deficits across a plethora of disorders.

## Introduction

Self-ordered sequencing is the ability to organize information and plan and execute a series of related steps in order to achieve a goal (Conen and Desrochers, 2022). Self-ordered sequencing is an executive cognitive function requiring elements of working memory, planning, and monitoring, occurring in a goal-directed fashion. The ability to self-order non-spatial sequences is disrupted by lesions to frontal and temporal lobes in humans (Petrides and Milner, 1982) and by mid-dorsolateral prefrontal cortex (PFC) lesions in rhesus monkeys (Petrides, 1995). Spatial working memory deficits in tasks requiring planning of self-ordered response sequences are also observed in both humans and rhesus monkeys following damage to the dorsolateral (dl)PFC (Owen et al., 1990; Passingham, 1985). Spatial sequencing deficits are also present in people with disorders such as obsessive-compulsive disorder (Banca et al., 2023; Vaghi et al., 2017), schizophrenia (Pantelis et al., 1997), Alzheimer’s (Lafleche and Albert, 1995), and Parkinson’s (Cooper et al., 1991) diseases. In the marmoset monkey, fiber-sparing lesions to the combined lateral, orbital, and anteroventral regions of the PFC confirmed its role in self-ordered sequencing (Collins et al., 1998). Subsequently, this effect was localized to ventrolateral (vl)PFC, but not orbitofrontal cortex (Axelsson et al., 2021; Walker et al., 2009).

In the marmoset, vlPFC projects to the striatum via glutamatergic pyramidal neurons (Roberts et al., 2007) as a relay of a fronto-striatal ‘loop’ (Alexander et al., 1986). The striatum is also implicated in self-ordered sequencing in humans (Jahanshahi et al., 2002) with potential contributions from both the caudate and putamen, which are implicated respectively in flexible goal-directed behavior and relatively automatic stimulus-response habits (Balleine and O’Doherty, 2010; Grahn et al., 2008). Previously, we have shown that the contribution of vlPFC in marmosets to self-ordered response sequencing of spatial arrays is limited to when the arrays vary across trials, requiring flexible, goal-directed behavior for successful performance (Axelsson et al., 2021). In contrast, when the same spatial array was used across trials in a fixed version of the self-ordered sequencing task which encouraged more automated responding involving use of a restricted set of preferred response sequences, inactivation of the vlPFC was without effect (Axelsson et al., 2021).

In the variable spatial array version of the self-ordered sequencing task, three stimuli appear in a random position on a computer screen for each trial and the animal must press each stimulus once and once only without returning to a previously selected one. Thus, in this version of the task, the animal must generalize application of a rule across trials with differing arrays, a process which involves executive functions including working memory and flexible planning. In contrast, for the fixed array, stimuli invariably appear in the same three locations. However, the marmoset can select whichever sequence of responding in these locations it prefers, thus allowing the animal to optimize its performance by utilizing more automated response sequences. Variable array responding is hypothetically more goal-directed (i.e., flexible) than fixed array responding, which is more habitual (i.e., automatic and rigid) (Dezfouli et al., 2014). Impairments in both forms of sequencing could arise from a tendency to maladaptively repeat responding (perseveration), a tendency that can arise in different forms (Sandson and Albert, 1984). Thus, repeating the just-performed response is termed continuous perseveration and interference caused by repetition later in the sequence is termed recurrent perseveration.

One aim of this study was to determine whether the caudate nucleus and/or putamen are causally implicated in spatial self-ordered sequencing behavior previously associated with upstream vlPFC function. The second aim was to determine whether the caudate and putamen have differential contributions to self-ordered sequencing, based on their respective roles in goal-directed and habitual behavior. To address these aims, we blocked glutamatergic transmission from cortical inputs to the striatum by microinfusions of AMPA (α-amino-3-hydroxy-5-methyl-4-isoxazole-propionate)/kainate glutamate receptor competitive antagonist CNQX (6-cyano-7-nitroquinoxaline-2,3-dione) (Smith et al., 1996) into those regions of caudate and putamen in receipt of projections from the vlPFC.

## Methods

### Subjects

Six common marmosets (*Callithrix jacchus*; see Supplemental Table 1A), were bred on-site at the University of Cambridge marmoset breeding colony. The marmoset holding rooms were kept at a constant 24°C with relative humidity of 55%. Holding rooms were gradually illuminated from 7:30 to 8 A.M. and gradually dimmed from 7:30 to 8 P.M., for a 12/12 h light/dark cycle with 30 min of dusk/dawn. Cages (2.8 × 1.2 × 0.98 m) contained a food tray, a nest box, wooden platforms at different heights and a variety of enrichment objects, including ladders, wooden branches, and ropes. Five days a week, animals had access to water for 2 h after behavioral testing and during this time period were fed MP.E1 primate diet (Special Diet Services) and carrots. During weekends animals had *ad libitum* access to water and were fed a calorically equal diet consisting of bread, egg, rusk, fruit and nuts. All procedures were conducted in accordance with the UK Animals (Scientific Procedures) Act 1986 as amended in 2012, under project license P09631465. In addition, the University of Cambridge Animal Welfare and Ethical Review Body (AWERB) provided ethical approval of the project license and its amendments, as well as individual studies and procedures via delegation of authorization to the NACWO for individual study plans. The study is summarized in Figure 1A.

**Figure 1:**
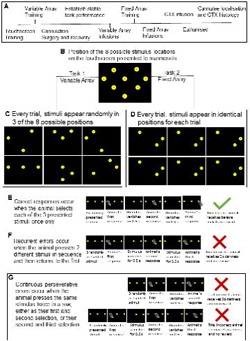
Outline of the spatial self-ordered sequencing task showing the experimental timeline (A), touchscreen stimuli positions (B), variable and fixed array tasks (C-D). An example of a correct sequence is shown (E), as well as perseverative errors of the recurrent (F) and continuous (G) type.

### Apparatus

All behavioral testing was performed in a custom-built testing apparatus located in a separate room from the marmoset holding rooms. Animals were trained to enter a custom-made Perspex transport box (Biotronix), in which they remained during testing. A door on the box was removed to provide access to a touch sensitive computer monitor (NEX121 TFT LCD Monitor, Nexio). Animals had to reach through an array of vertical bars to respond to visual stimuli on the touch screen. Reward, in the form of banana milk (Nesquik banana powder in milk, Nestlé), was delivered through a peristaltic pump to a licking spout with lick sensor which was accessible through the vertical bars. Auditory stimuli were presented through a speaker, out of sight of the subjects. Reward delivery and presentation of visual and auditory stimuli were controlled by the application MonkeyCantab (R. N. Cardinal), using the Whisker control system (Cardinal and Aitken, 2010).

### Behavioral Training

#### Pre-operative Training

Subjects were trained to enter the transport box and habituated to the test apparatus. After successful habituation, animals were familiarized with the liquid reward, learned the association between an auditory stimulus and access to reward and then acquired a touchscreen response for that reward; all previously described in Roberts et al., (1988). Subsequently, animals were trained on a spatial self-ordered sequencing task (Figure 1B) designed in Monkey Cantab by AC Roberts and TW Robbins, in which they were required to select each of an array of identical stimuli presented on the screen, once only. Subjects were first trained to touch a stimulus, presented in a random spatial location for each trial, based on an eight-location grid. Once animals performed 20 trials in a session, task difficulty was gradually increased. The first step of subsequent training was the addition of a second identical stimulus in a distinct spatial location and animals were required to respond to both spatial locations, sequentially, in any order, to receive reward. Once a response was made to a stimulus, that stimulus disappeared for a set amount of time, denoted “vanishing time” (vt). Animals were allowed to continue responding throughout the vt, but if they responded to the same stimulus more than once, the trial ended prematurely, the houselight was turned off for 5 s and the trial scored as incorrect. Vt was gradually decreased during training and the number of stimuli were increased until animals could perform 20 two-stimuli trials with an accuracy of 80%, then 30-40 three-stimuli trials with an accuracy equal to or greater than 50%, both with a vt = 0.5 s.

#### Post-operative training

After surgery, a stable baseline was established in animals on the final version of the variable array, with the most difficult parameters; three visual stimuli with a vt of 0.5 s. When animals consistently performed above 50% accuracy and were habituated to their holders (experimental staff that would gently hold them for infusions) such that task accuracy was not impaired, they began receiving infusions into either the caudate nucleus, putamen, or combined caudate and putamen.

#### Task versions: variable array vs fixed array

Two versions of a spatial self-ordered sequencing task were used for these experiments involving either a variable array (Figure 1C), or a fixed array (Figure 1D), as used previously (Axelsson et al., 2021). In the variable array, stimuli appeared randomly in three of eight possible locations which varied across trials. In the fixed array, animals were presented with three stimuli that always appeared in the same locations across trials and sessions. The variable array required animals to flexibly create and apply a plan to action a correct self-ordered spatial sequence, thereby applying a generalized abstract schema to solve the problem. The fixed array allowed animals to develop an optimal response strategy (or sequence) to complete the task in a more habitual manner. Both tasks required subjects to self-order and execute a sequence of responses to three identical, but spatially distinct stimuli presented on a screen, touching each stimulus once and only once, with a vt of 0.5 s. If the animal touched the same stimulus twice within a trial – either twice consecutively (a continuous perseverative error) or returned to the first stimulus after selecting two different stimuli (a recurrent perseverative error) – the trial was deemed incorrect, terminated early, and the animal experienced 5 s of darkness before the next trial commenced.

Spatial locations of the three stimuli in the fixed array were selected specifically for each animal, thus whilst constant across trials and sessions for an individual animal, it varied between animals (Supplemental Table 1A). By presenting the same array to an animal repeatedly across trials and sessions it was hypothesized that they would develop a preferred sequence strategy across time to reduce working memory load. Having shown development of a response sequence strategy the effects of striatal manipulations would then be compared to that seen on variable arrays where there was a greater working memory load. Thus, the set of three stimuli chosen for each animal was based on their overall performance of this particular array during variable task 1. First, subjects had to have performed at least five of the six possible correct response sequences for the proposed array. Second, animals didn’t show a strong preference for any particular one of the response sequences (i.e., the Shannon entropy value was as close to that predicted if responses were random, 2.58 bits). Third, animals had an accuracy score for the specific fixed sequence of at least 50%, which was considerably superior to chance performance (22%).

#### Surgery

Animals had permanent indwelling cannulae implanted to allow infusion of drugs into the caudate nucleus, putamen, or both together. For surgery, animals were premedicated with 0.1 ml of 100 mg/ml ketamine (Ketavet, Henry Schein Medical) and given prophylactic analgesic (0.075 ml of 50 mg/ml subcutaneous metacam, Pfizer) before being intubated (using Intubeaze 20mg/ml lidocaine hydrochloride spray, Dechra Veterinary Products Ltd., Shropshire, UK) and anesthesia maintained using a mixture of vaporized isoflurane (Novartis Animal Health) and O_2_ (2.25% isoflurane in 0.3 l/min O_2_). Animals were then placed in a marmoset stereotaxic frame (David Kopf). Heart rate, O_2_ saturation, respiratory rate, and CO_2_ saturation were all monitored by pulse oximetry and capnography (Microcap Handheld Capnograph, Oridion Capnography Inc., MA, USA) while core body temperature was monitored rectally (TES-1319 K-type digital thermometer, TES Electrical Electronic Corp., Taipei, Taiwan).

Cortical depth was measured at +17.5 on the anterior-posterior (AP) axis, and -1.5 on the latero-medial (LM) axis to allow for corrections to cannula target placements, as previously described in Dias et al., (1997). Double guide cannulae measuring 7 mm in length with a center-center distance of 1.4 mm (Plastics One) were then inserted, with the medial guide at AP +11 LM +/-3.3. Precise locations of all cannulae in all animals were determined using post-mortem histology (Figure 2). Guides were fixed in place by skull screws and dental acrylic (Simplex Rapid, Kemdent Works). Post-surgically, subjects were administered 0.18 ml of 3.8 mg/ml dexamethasone (0.09 ml injected into each quadricep; Aspen Pharma Trading Ltd.). Subjects were also given analgesic once daily in the morning, for 3 d after surgery (meloxicam, 0.1 ml of a 1.5 mg/ml oral suspension; Boehringer Ingelheim). After surgery animals had *ad libitum* access to water for at least two weeks and were provided food that was otherwise only available to them on weekends.

**Figure 2:**
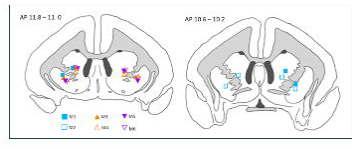
Cannulae placements for each of the six experimental marmosets located within the vlPFC target regions within the caudate and putamen. Males are open symbols, females filled symbols. Square, M1 and M2; Triangle, M3 and M4; Inverted triangle, M5 and M6.

**Figure 3:**
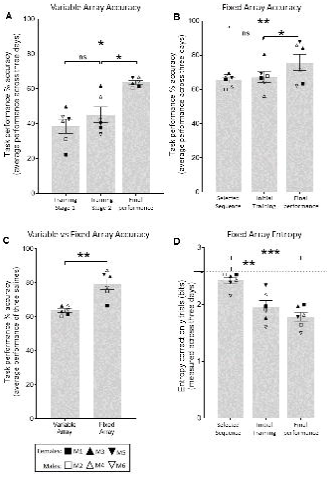
Task performance accuracy on variable and fixed array tasks (A, B, C), and entropy measures across training compared to saline infusion performance on the fixed task (D). Arrow on y axis indicates the entropy score if correct sequences were selected randomly (2.58 bits). For the variable array (A) one-way rmANOVA revealed a main effect of Training Stage on accuracy (F (2,10) = 20.45; p=0.0003); Sidak-corrected posthoc showed: Training Stage 1 vs Final Performance (p=0.0032), Training Stage 2 vs Final Performance (p=0.0003). For the fixed array (B) one-way rmANOVA showed a main effect of Training Stage on accuracy (F (2, 10) = 8.512; p=0.0069); Sidak-corrected posthoc showed: Selected Sequence vs Final Performance (p=0.0099), and Training Stage 1 vs Final Performance (p=0.0263). The significant difference between variable and fixed array accuracy (C) was determined via paired t-test (p=0.0014). Finally, a one-way rmANOVA showed a main effect of Training Stage on entropy (F (2,10) = 20.81; p=0.0003); Sidak-corrected posthoc showed: Selected Sequence vs Initial Training (p=0.0044), Selected Sequence vs Final Performance (p=0.0003). Data are represented as mean ± standard error of mean and Sidak comparisons indicated by *p>0.05, **p>0.01, and ***p>0.001; ns=not significant. Data are represented as mean ± standard error of mean. Data were collected from 6 monkeys, each with their own symbol as designated in Figure 2.

#### Drug preparation and treatment

For drug infusions, animals were gently restrained by a familiar person (holder) other than the researcher and taken to a designated infusion-room. The researcher gently removed caps and dummies from cannula guides and cleaned the guides with injection wipes. For all infusions, an injector (Plastics One) was used that protruded +0.1 mm from the cannula directed to the caudate nucleus or +0.3 mm from the cannula directed to the putamen. The injectors were connected to a 10 µl Hamilton syringe (701RN; Hamilton) via PTFE tubing (0.3 mm in diameter). Solvent flexible tubing was used to connect PFTE tubing to injector and syringe (0.38 mm in inner diameter, Elkay Laboratory Products, Ltd.). Drug was accurately delivered by an infusion pump (KDS230, KD Scientific). Injectors, tubing and syringes were all sterilized before setup.

CNQX disodium salt (Tocris; MW 312.15), an AMPA receptor antagonist, was dissolved in sterile saline (Hameln) to a 3 mM solution, and further diluted in sterile saline to 1 mM solution. Solutions were filtered (Whatman Uniflo syringe filter; 0.2 µm pore; Cytiva) and aliquoted into sterile eppendorfs for immediate use or storage at -20 °C for up to one month. After one month, new solutions were made. For control infusions, sterile saline was infused instead of CNQX.

#### Experimental design, measurements, and statistical analysis

Animals performed the task once daily on Monday through to Friday every week, at approximately the same time each day. A within-subject study design was used whereby each animal performed both tasks and received all control and drug infusions across all brain regions. All animals started on the more difficult variable array (Task 1) and moved to the fixed array (Task 2) with some animals returning to variable after being on fixed (detailed in Supplemental Table 1B). Drug dose and brain region were counterbalanced across all animals, as summarized in Supplemental Table1C-F. Typically, animals received a mock infusion on Wednesday, saline infusion on Thursday, and a CNQX infusion on Friday. Occasionally, animals received a mock on Tuesday, saline infusion on Wednesday, and CNQX on Thursday. A paired t-test revealed no significant difference between saline control infusions for 1 mM and 3 mM CNQX, and thus control performance was averaged for each animal for each brain region for comparisons to 1 mM and 3 mM CNQX.

Before infusions began, animals were required to reach stable performance such that it was reliable across the proposed infusion days and was not affected by handling of the animal prior to sessions with variable (Supplemental Figure 1) and fixed (Supplemental Figure 2) arrays. Animals received acute infusions of low concentration (1 mM) or high concentration (3 mM) CNQX ((6-cyano-7-nitroquinoxaline-2,3-dione), an AMPA/kainate glutamate receptor competitive antagonist) or saline into either the caudate nucleus, putamen, or both together. Since most of the input into the striatum is glutamatergic, CNQX infusions acted to inhibit that input. The effects of these infusions on task performance were investigated.

Response variables measured include the number of trials completed, accuracy, and type and number of errors performed, and omissions. Trials completed were the total number of trials performed, including correct, incorrect, and omissions. If three consecutive omissions were made (i.e., the animal did not respond to stimuli presented for 60 s on three consecutive occasions), the session ended.

Accuracy was the number of sequences performed correctly divided by the number of trials completed minus the number of omissions (correct trials / (total trials – omissions)). Errors per session were grouped together based on whether they were continuous or recurrent. Error types were presented as a percentage of total trials (continuous errors / (total trials – omissions); recurrent errors / (total trials – omissions)).

Entropy was calculated for the fixed array after selecting only correct responses and analyzing the proportion of each of the six possible correct patterns employed by the animals using the following equation:

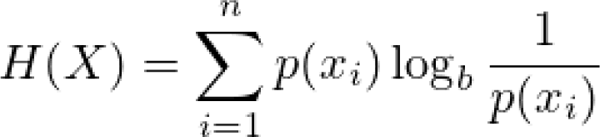

Testing data were collected in a Microsoft Access database. Data were extracted and organized, using R studio (version 4.1.2 (2021-11-01), Rstudio: Integrated Development for R; Rstudio) and Microsoft Excel (Office 16). Statistical analysis and graphical representation were performed in GraphPad Prism (version 9.3.1 for Windows, GraphPad Software). Data are presented as mean values ± SEM. One-way or two-way repeated measures ANOVAs were performed on data followed by post-hoc corrections for multiple comparisons, as appropriate and indicated in Figure legends.. Results were considered significant when *p* < 0.05. For all data displayed in Figures 4 and 5, a two-way ANOVA with factors of Treatment (Saline, 1 mM CNQX, 3 mM CNQX) and Brain Region (caudate nucleus, putamen, or combined caudate and putamen) were employed, followed by Šídák’s multiple comparisons test.

**Figure 4:**
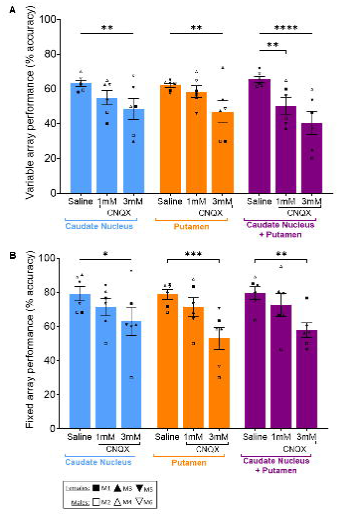
Effects of 1 mM or 3 mM CNQX into either caudate nucleus, putamen, or combined caudate and putamen on variable (A) and fixed (B) array accuracy. Data were collected across all regions and both CNQX doses from 6 monkeys, each with their own symbol defined in the key. For the variable array (A) two-way repeated measures (rm)ANOVA revealed a main effect of Treatment (F (2, 10) = 13.04; p=0.0016); but no effect of brain region (F<1). Sidak-corrected posthoc showed: caudate saline vs 3 mM CNQX (p=0.0088), putamen saline vs 3 mM CNQX (p=0.0073), combined caudate+putamen saline vs 1 mM or 3 mM CNQX (p=0.0064, p<0.0001, respectively). The fixed array (B), two-way rmANOVA revealed a main effect of Treatment (F (2, 10) = 23.91; p=0.0002) but no effect of brain region F (2, 10) = 2.495; p=0.13211); Sidak-corrected posthoc showed: caudate saline vs 3 mM CNQX (p=0.265), putamen saline vs 3 mM CNQX (p=0.0007), combined caudate+putamen saline vs 3 mM CNQX (p=0.0033,). Data are represented as mean ± standard error of mean and Sidak comparisons indicated by *p>0.05, **p>0.01, ***p>0.001; or ***p>0.0001.

**Figure 5:**
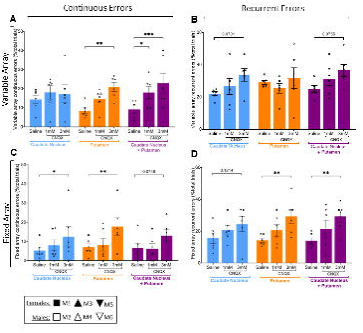
Effects of 1 mM or 3 mM CNQX into either caudate nucleus, putamen, or combined caudate and putamen on perseverative errors of the continuous (A,C) and recurrent type (B,D) in variable (A,B) and fixed (C,D) arrays. Data were collected across all timepoints from 6 monkeys, each with their own symbol defined in the key. For the variable array two-way rmANOVAs revealed a main effect of Treatment on continuous (F (2, 10) = 4.694; p=0.0365) and recurrent errors (F (2, 10) = 7.287; p=0.0112) but no effect on brain region (F<1 for both continuous and recurrent errors). Sidak-corrected posthoc showed for variable array continuous errors: putamen saline vs 3 mM CNQX (p=0.0011), and combined caudate+putamen saline vs 1 mM or 3 mM CNQX (p=0.0164, p=0.0004, respectively). For variable array recurrent errors, it revealed: caudate saline vs 3 mM CNQX (p=0.0701) and combined caudate+putamen saline vs 3 mM CNQX (p=0.0755). The fixed array two-way rmANOVAs revealed a main effect of Treatment on both continuous (F (2, 10) = 6.481; p=0.0157) and recurrent errors (F (2, 10) = 15.56; p=0.0009) but no effect of brain region ( F2, 10) = 2.477; p=0.1337 for continuous errors, and F<1 for recurrent errors). Sidak-corrected posthoc showed for fixed array continuous errors: caudate saline vs 3 mM CNQX (p=0.0429), putamen saline vs 3 mM CNQX (p=0.0029). For fixed array recurrent errors, it revealed: putamen saline vs 3 mM CNQX (p=0.0069), combined caudate+putamen saline vs 3 mM CNQX (p=0.0062). Data are represented as mean ± standard error of mean and Sidak comparisons indicated by *p>0.05,**p>0.01, or ***p>0.001.

## Results

### Accuracy on the fixed array task was greater than that of the variable array task, a likely consequence of increased automaticity of responding

Marmoset accuracy on the variable array improved significantly after training when comparing initial, to final, task performance after control saline infusions (Figure 3A). This performance then reached asymptote, remaining stable for the duration of the variable array component of the study, as confirmed by a one-way rmANOVA which revealed no main effect across time on accuracy of performance measured across all saline control infusions (F (5, 25) = 0.3756, p=0.8606; data not shown). Overall performance improved after transfer from variable to fixed array (Figure 3B), where animals reached asymptote at a significantly higher level for the fixed array (Figure 3C). This increased accuracy was associated with a decrease in entropy (Figure 3D) which occurred because marmosets developed a preferred set of ‘correct’ response sequences out of the 6 possible ‘correct’ sequencies available (Supplemental Table 2A-B). Development of a preferred pattern of responding is consistent with the animals’ performance of the fixed array task becoming increasingly automated. Post-training, accuracy remained stable for the duration of the fixed array component of the study, as determined by a one-way rmANOVA which revealed no main effect of time on accuracy of performance across saline control infusions (F (5, 25) = 1.808, p=0.1477; data not shown).

### High dose CNQX (3 mM) impaired accuracy on both variable and fixed array tasks in the caudate nucleus, putamen, or both combined

High dose CNQX (3 mM) significantly impaired accuracy in both variable and fixed arrays regardless of striatal location (Figure 4). On the variable array (Figure 4A), 3 mM CNQX impaired accuracy to similar extents when infused into either the caudate or putamen alone. However, when combined, the impairments were additive, leading to a greater decline in accuracy than when either caudate or putamen received 3 mM CNQX alone. In the fixed array (Figure 4B), these same manipulations caused a decline in accuracy, but the effect appeared greatest when the putamen was targeted alone.

### High dose CNQX (3 mM) had differing effects on error types in both variable and fixed arrays depending on brain region(s) targeted

For the variable array (Figure 5A-B), there was an increase in ‘continuous’, but not ‘recurrent’ errors after CNQX into the putamen. In contrast, there was no effect on continuous errors after CNQX into the caudate (Figure 5A), though recurrent errors trended towards an increase (p=0.0701; Figure 5B). Continuous errors significantly increased when the caudate and putamen simultaneously received 3 mM CNQX (Figure 5A), which resembled the effect seen when the putamen alone was targeted. Similarly, there was a trend towards increased recurrent errors after combined targeting of caudate and putamen (p=0.0755; Figure 5B), comparable to when the caudate was targeted alone.

In the fixed array (Figure 5C-D), continuous errors increased significantly after 3 mM CNQX into the caudate, though recurrent errors were unaffected. However, both continuous and recurrent errors increased after 3 mM CNQX into the putamen. Whilst there was only a trend (p=0.0748) towards increased continuous errors after 3 mM CNQX infusion into caudate and putamen combined (Figure 5C), there was a significant increase in recurrent errors (Figure 5D).

### Low dose CNQX (1 mM) only impaired accuracy on variable arrays following combined infusions into the caudate and putamen

Low dose CNQX (1 mM) into either caudate nucleus or putamen individually did not impair accuracy for either variable or fixed arrays (Figure 4). However, for the variable array (Figure 4A), when both caudate nucleus and putamen received 1 mM CNQX simultaneously, accuracy was significantly impaired, associated with an increase in continuous errors (Figure 5A). In contrast, the same combined manipulation did not affect accuracy in the fixed array (Figure 4B).

The main experimental findings from this study are summarized in Table 1

**Table 1:**
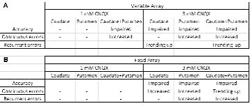
Summary of effects seen after infusions on the variable (A) and fixed arrays (B)

## Discussion

Previously, inactivation of the vlPFC impaired spatial self-ordered sequencing in the variable, but not fixed array (Axelsson et al., 2021). In this study, only low dose CNQX, targeting vlPFC downstream glutamatergic striatal inputs, selectively impaired variable array performance and only when infused into caudate and putamen combined. In contrast, we generally found impairments in both tasks with high dose CNQX, which impaired accuracy on both variable and fixed arrays in caudate, putamen, or both combined. There were differing effects on error types though, dependent on striatal region and whether the array was variable or fixed.

Specifically, CNQX into putamen, but not caudate, selectively enhanced continuous perseverative responses in the variable array, which requires the animal to use an overall schema, plan or strategy to guide performance. Instead, caudate infusions selectively increased recurrent perseverative errors in the same array. These regionally differential effects in the case of the putamen could be attributed to response sequencing deficits resulting from a failure to monitor motor feedback from the immediately preceding response, thus resulting in motor perseveration. For the caudate nucleus, increased recurrent errors likely reflect interference in spatial working memory, broadly consistent with predicted roles of the caudate nucleus in working memory (Akhlaghpour et al., 2016; Collins et al., 2000; Grahn et al., 2008; Kermadi and Joseph, 1995; Levy et al., 1997; Lewis et al., 2004; Postle and D’Esposito, 1999) and the putamen in motor function (Kubota et al., 2009; Turner et al., 2022; Yin, 2010; Yin et al., 2009).

A greater role for the striatum than the vlPFC in controlling responding in the fixed array was hypothesized since the putamen, in particular, has been implicated in learning and performance of skilled or habitual motor sequences in rodents (Kubota et al., 2009; Turner et al., 2022; Yin, 2010; Yin et al., 2009). In the present study, we demonstrate development of greater response automaticity in the fixed array as evidenced by a decrease in entropy, suggesting skilled or habitual control (Dezfouli et al., 2014). Consistent with the rodent evidence there was some indication here that glutamate receptor blockade in the putamen produced deficits that were greater than in the caudate for the fixed array task and comparable with combined infusions into the caudate and putamen (Fig 4B). Moreover, the effects of putamen infusions on continuous errors were greater than for either the caudate, or combined infusions (Fig 5C). However, it should be noted that there were evidently some marginal detrimental effects of caudate CNQX infusions on fixed array performance and we did not demonstrate greater selectivity in the putamen at the lower dose of CNQX.

For the variable array, both caudate and putamen infusions impaired performance, with combined infusions being additive even at low dose (as well as high dose) CNQX; unlike for the fixed array (Fig 4) where effects were not additive at either dose. This is consistent with evidence that the vlPFC projects to both striatal regions targeted by these infusions (Roberts et al., 2007) which may suggest a combined role in flexible responding. Indeed, a human fMRI study demonstrates that anterior putamen (in addition to caudate) is active during the planning stage of variable responses to numerical sequences (Jankowski et al., 2009). However, given that the caudate and putamen have opposing roles in reversal learning in marmoset (Jackson et al., 2019) and in response sequencing in rat (Turner et al., 2022), it is also plausible that the two regions mediate distinct aspects of performance. In the present study, the pattern of perseverative errors differed for the two sites; putamen being associated with elevated continuous errors, and caudate being associated with increased recurrent errors. This is consistent with inactivation of dorsolateral striatum (i.e., rodent putamen) inducing continuous errors in a serial order task (Rothwell et al., 2015), and a role of dorsomedial striatum (i.e., rodent caudate) in self-monitoring during execution of goal-directed action sequences (Vandaele et al., 2021). Such a pattern implies deficits in spatial working memory and impaired motor control, respectively (see above), although it should be noted that both tasks were designed to minimize the load of holding online motor sequences in working memory (Collins et al., 1998). A more general strategic deficit in applying a strategy or schema to the sequencing task (Owen et al., 1990) might be implicated instead. Inactivation of the vlPFC or antagonism of its 5-HT2A receptors increased only recurrent errors in variable arrays, although dopamine D2 receptor antagonism did enhance continuous perseverative errors (Axelsson et al., 2021). Hence, it is possible that different projections of the vlPFC to the striatum contribute to different aspects of sequencing performance in the variable condition. Of course, it is also plausible that influence of other cortical or thalamic projections to these regions of the striatum was impaired by intra-striatal blockade of glutamate receptors. Indeed, the lack of effect of vlPFC infusions on fixed array performance suggests that striatal involvement in this task is mediated by different cortical inputs, hypothetically involving cortical sensorimotor regions (Sánchez-Fuentes et al., 2021).

The striatum receives vast numbers of glutamatergic inputs from cortical and subcortical regions which converge anatomically with dopaminergic afferents from the midbrain. All major glutamate ionotropic (NMDA (N-methyl-D-aspartate), kainate, and AMPA) as well as metabotropic receptors are represented. AMPA-receptors in the striatum are located on terminals of corticostriatal afferents (Fujiyama et al., 2004), and act to regulate glutamate release through a positive feedback mechanism (Fujiyama et al., 2004; Patel et al., 2001). To this effect, 1 mM CNQX has successfully been used in the dorsal striatum of rats to alter the balance between goal-directed and habitual control of behavior (Furlong et al., 2014), as well as altering control of goal-directed behavior following intra-caudate infusions in marmosets (Duan et al., 2021). Here, a possible limitation is that main effects were only observed after 3 mM CNQX infusions. Whilst CNQX is relatively selective for AMPA receptors (IC value 400 nM) with additional actions on kainate receptors (IC, 4 µM), and at high concentrations, NMDA receptors (Brickley et al., 2001; Lester et al., 1988; Stein and Verdoorn, 1992) it is likely that, at concentrations used here, CNQX non-selectively impaired striatal glutamatergic transmission.

## Conclusions

By employing local intra-striatal CNQX infusions to block glutamatergic transmission, we implicate marmoset lateral prefrontal cortex output striatal regions (caudate and putamen) in self-ordered sequencing behavior. Both flexible responding driven by caudate, and more rigid habitual responding driven by putamen are differentially required for marmosets to successfully self-order a sequence in variable and fixed arrays. Caudate and putamen have additive roles in variable (but not fixed) array self-ordered sequencing, and infusions into the two regions induced different kinds of perseverative errors. By understanding the basic neural circuitry underlying such sequencing abilities, we may better understand how to improve them in people with deficits associated with neuropsychiatric disorders, such as obsessive-compulsive disorder, as well as basal ganglia disorders.

