## Supplementary figures and images for "Comparative roles of caudate and putamen in the serial order of behavior: Effects of striatal glutamate receptor blockade on variable versus fixed spatial self-ordered sequencing in marmosets"

### Supplemental Figure 1

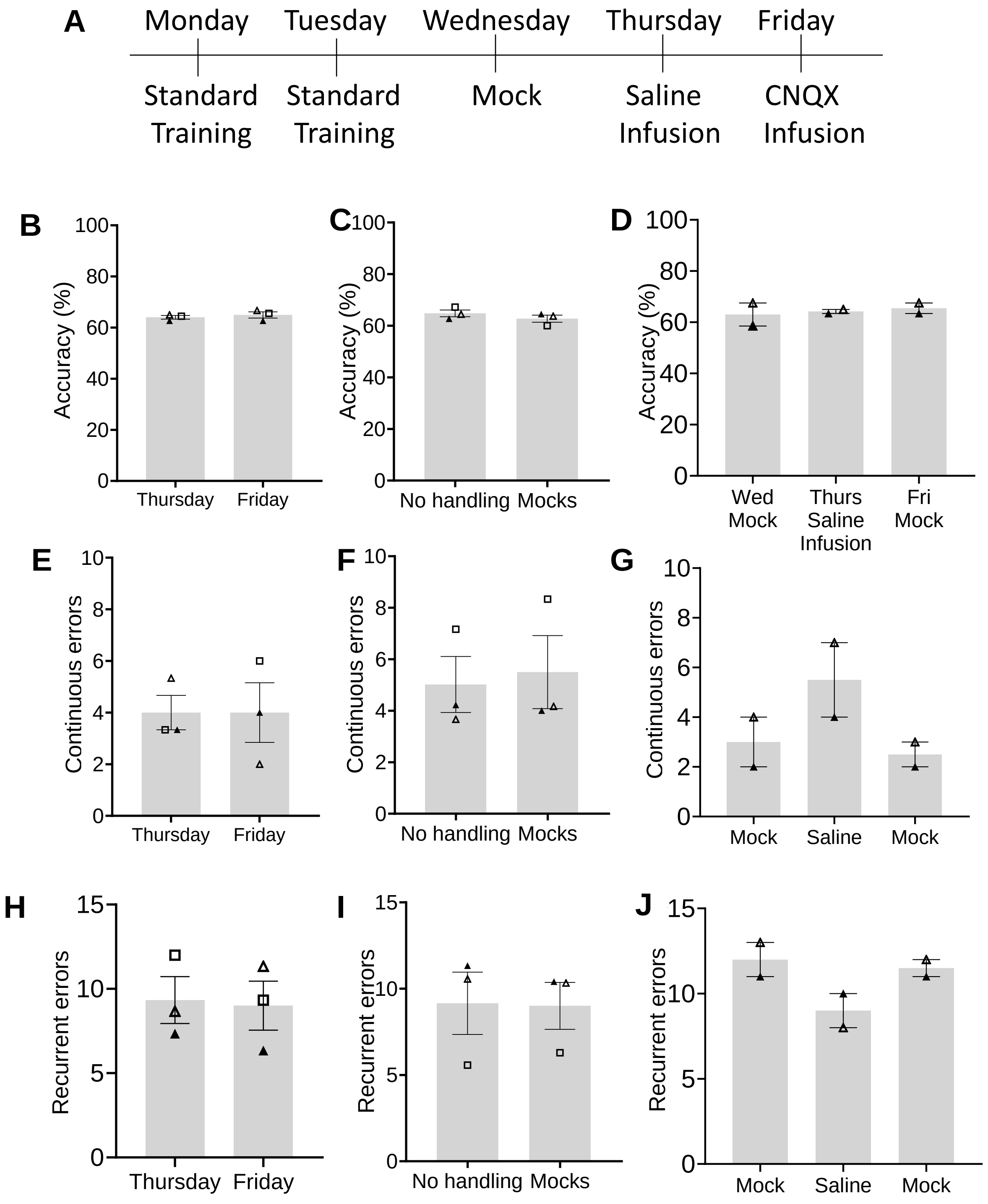

### Supplemental Figure 2

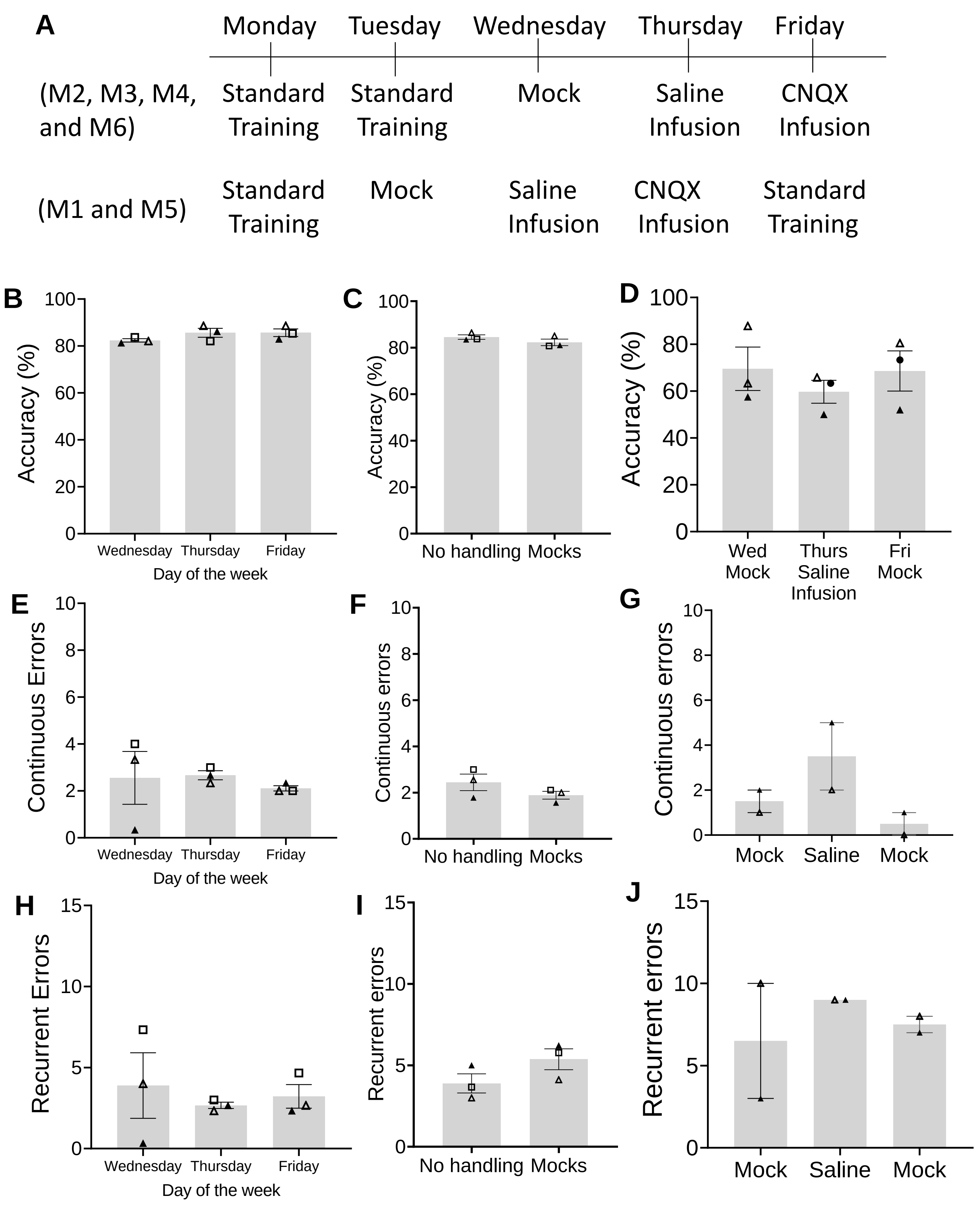
