## Supplemental Table 1 for "Comparative roles of caudate and putamen in the serial order of behavior: Effects of striatal glutamate receptor blockade on variable versus fixed spatial self-ordered sequencing in marmosets"

**A**

| Animal | Symbol | Sex | Fixed Array Pattern | Infusion type performed | | | Total Infusions | | Tract tracing infusion |
|---|---|---|---|---|---|---|---|---|---|
| | | | | Caudate | Putamen | C+P | Caudate | Putamen | |
| M1 | ■ | Female | 642 | 15 | 14 | 13 | 28 | 27 | N/A |
| M2 | □ | Male | 750 | 16 | 16 | 14 | 30 | 30 | N/A |
| M3 | ▲ | Female | 631 | 12 | 15 | 11 | 23 | 26 | CTX (left caudate) |
| M4 | △ | Male | 621 | 9 | 10 | 9 | 18 | 19 | CTX (left Putamen) |
| M5 | ▼ | Female | 540 | 8 | 8 | 12 | 20 | 20 | CTX (left Putamen) |
| M6 | ▽ | Male | 765 | 12 | 12 | 11 | 23 | 23 | CTX (left caudate) |

**B**

| **CNQX Concentration and Array Type** | | **Animal** | | | | | |
|---|---|---|---|---|---|---|---|
| | | **M1** | **M2** | **M3** | **M4** | **M5** | **M6** |
| **Dose/ Array** | **1 mM Variable** | First | First | First | First | Second | First |
| | **3 mM Variable** | Fourth | Third | Second | Second | First | Second |
| | **1 mM Fixed** | Second | Second | Third | Third | Fourth | Fourth |
| | **3 mM Fixed** | Third | Fourth | Fourth | Fourth | Third | Third |

**C**

| **Variable Array 1 mM CNQX** | | **Animal** | | | | | |
|---|---|---|---|---|---|---|---|
| | | **M1** | **M2** | **M3** | **M4** | **M5** | **M6** |
| **Brain Region** | **Caudate** | First | First | Third | Third | Third | Second |
| | **Putamen** | Second | Second | First | First | Second | Third |
| | **Caudate+Putamen** | Third | Third | Second | Second | First | First |

**D**

| **Variable Array 3 mM CNQX** | | **Animal** | | | | | |
|---|---|---|---|---|---|---|---|
| | | **M1** | **M2** | **M3** | **M4** | **M5** | **M6** |
| **Brain Region** | **Caudate** | Second | Second | Third | Third | Third | First |
| | **Putamen** | First | Third | First | First | Second | Second |
| | **Caudate+Putamen** | Third | First | Second | Second | First | Third |

**E**

| **Fixed Array 1 mM CNQX** | | **Animal** | | | | | |
|---|---|---|---|---|---|---|---|
| | | **M1** | **M2** | **M3** | **M4** | **M5** | **M6** |
| **Brain Region** | **Caudate** | First | First | Third | Third | Second | Second |
| | **Putamen** | Second | Second | First | First | First | First |
| | **Caudate+Putamen** | Third | Third | Second | Second | Third | Third |

**F**

| **Fixed Array 3 mM CNQX** | | **Animal** | | | | | |
|---|---|---|---|---|---|---|---|
| | | **M1** | **M2** | **M3** | **M4** | **M5** | **M6** |
| **Brain Region** | **Caudate** | Second | Second | Third | Third | Third | First |
| | **Putamen** | First | Third | First | First | Second | Second |
| | **Caudate+Putamen** | Third | First | Second | Second | First | Third |
