## Supplemental Table 2 for "Comparative roles of caudate and putamen in the serial order of behavior: Effects of striatal glutamate receptor blockade on variable versus fixed spatial self-ordered sequencing in marmosets"

**A**

| Animal | Sequence | Trial Type | Number of different response patterns selected | | | | | | | | | | | |
|---|---|---|---|---|---|---|---|---|---|---|---|---|---|---|
| | | | Selected Sequence | First 5 days training | Average 5 baselines pre-infusions | Mocks | Caudate | | | Putamen | | | Caudate + Putamen | | |
| | | | | | | | Saline | 1 mM CNQX | 3 mM CNQX | Saline | 1 mM CNQX | 3 mM CNQX | Saline | 1 mM CNQX | 3 mM CNQX |
| M1 | 642 | Correct | 6.0 | 4.8 | 4.2 | 4.8 | 4.5 | 5.0 | 5.0 | 4.5 | 5.0 | 5.0 | 4.0 | 6.0 | 4.0 |
| M2 | 750 | Correct | 6.0 | 4.0 | 4.6 | 3.9 | 4.0 | 5.0 | 5.0 | 3.5 | 4.0 | 3.0 | 5.0 | 5.0 | 4.0 |
| M3 | 631 | Correct | 6.0 | 5.0 | 4.8 | 4.3 | 5.0 | 4.0 | 4.0 | 5.5 | 5.0 | 5.0 | 5.5 | 5.0 | 6.0 |
| M4 | 621 | Correct | 6.0 | 5.0 | 5.0 | 4.0 | 4.0 | 6.0 | 3.0 | 4.0 | 5.0 | 5.0 | 4.0 | 5.0 | 3.0 |
| M5 | 540 | Correct | 6.0 | 5.0 | 5.0 | 5.0 | 4.5 | 5.0 | 4.0 | 5.0 | 6.0 | 4.0 | 5.5 | 4.0 | 5.0 |
| M6 | 765 | Correct | 5.0 | 4.0 | 4.0 | 4.5 | 5.0 | 4.0 | 5.0 | 4.5 | 4.0 | 5.0 | 4.0 | 4.0 | 5.0 |
| **Average** | | **Correct** | **5.8** | **4.6** | **4.6** | **4.4** | **4.5** | **4.8** | **4.3** | **4.5** | **4.8** | **4.5** | **4.7** | **4.8** | **4.5** |
| **Median** | | **Correct** | **6.0** | **4.9** | **4.7** | **4.4** | **4.5** | **5.0** | **4.5** | **4.5** | **5.0** | **5.0** | **4.5** | **5.0** | **4.5** |

**B**

| Animal | Sequence | Trial Type | Number of different response patterns selected | | | | | | | | | | | |
|---|---|---|---|---|---|---|---|---|---|---|---|---|---|---|
| | | | Selected Sequence | First 5 days training | Average 5 baselines pre-infusions | Mocks | Caudate | | | Putamen | | | Caudate + Putamen | | |
| | | | | | | | Saline | 1 mM CNQX | 3 mM CNQX | Saline | 1 mM CNQX | 3 mM CNQX | Saline | 1 mM CNQX | 3 mM CNQX |
| M1 | 642 | All | 12.0 | 10.2 | 9.4 | 9.3 | 10.0 | 9.0 | 14.0 | 9.5 | 11.0 | 10.0 | 9.0 | 12.0 | 7.0 |
| M2 | 750 | All | 12.0 | 8.6 | 7.6 | 8.4 | 8.5 | 8.0 | 7.0 | 7.5 | 7.0 | 10.0 | 8.0 | 10.0 | 7.0 |
| M3 | 631 | All | 11.0 | 9.0 | 9.6 | 9.0 | 8.0 | 8.5 | 11.0 | 8.0 | 11.0 | 13.0 | 9.0 | 10.0 | 14.0 |
| M4 | 621 | All | 12.0 | 10.8 | 8.8 | 7.3 | 7.0 | 10.0 | 5.0 | 8.0 | 9.0 | 8.0 | 6.5 | 14.0 | 5.0 |
| M5 | 540 | All | 12.0 | 10.6 | 10.0 | 9.5 | 8.0 | 13.0 | 8.0 | 11.0 | 11.0 | 10.0 | 10.5 | 9.0 | 13.0 |
| M6 | 765 | All | 11.0 | 11.4 | 10.5 | 8.5 | 11.5 | 9.0 | 12.0 | 8.5 | 9.0 | 12.0 | 9.0 | 8.0 | 9.0 |
| **Average** | | **All** | **11.7** | **10.1** | **9.3** | **8.7** | **8.8** | **9.6** | **9.5** | **8.8** | **9.7** | **10.5** | **8.7** | **10.5** | **9.2** |
| **Median** | | **All** | **12.0** | **10.4** | **9.5** | **8.8** | **8.3** | **9.0** | **9.5** | **8.3** | **10.0** | **10.0** | **9.0** | **10.0** | **8.0** |
